# Strategic Conservation Prioritization Through Creating and Updating the Regional Species of Greatest Conservation Need (RSGCN) List in the Northeastern U.S.

**DOI:** 10.1101/2025.01.03.631241

**Authors:** Melissa D. Starking, Tracy Monegan Rice, Karen Terwilliger, Donovan Drummey, Mack Frantz, Catherine D. Haffner, Brian Hess, Lisa Holst, Michael Marchand, Kevin R. Burgio

## Abstract

Through the end of the century, biodiversity is expected to markedly decline around the world due to climate change, habitat loss, and other factors. Because of this, there is an increasing need for more efficient, effective, and collaborative conservation efforts worldwide. While many global and national threat assessment databases exist, they may only have limited utility at regional and local scales where on-the-ground conservation is best implemented. To better identify species risk at a more regional scale, the 14 state fish and wildlife agencies in the northeastern United States worked with taxonomic experts across the Northeast to create the most recent (2023) list of Regional Species of Greatest Conservation Need (RSGCN). We compared a preliminary list compiled from global and national datasets to the finalized list after the process documented here. We found that the global and national datasets only identified ∼55% of the 806 species statuses accurately for the region, demonstrating the importance of current local and regional expertise and data. The process of identifying RSGCN for the northeastern U.S. has resulted in specific actionable information leading to beneficial outcomes, such as dedicated funding for regional initiatives, data-sharing, and coordination among regional, state, and local conservation organizations - enabling beneficial conservation outcomes across the region for many of the species and habitats of the region. The methods described may serve as a framework for other regions to help identify conservation targets using the best available and more localized landscape- and watershed-scale information.

## Introduction

Due to the predicted acceleration of extinction events due to climate change and habitat loss (e.g., Urban 2015, Carlson et al. 2017, Sandor et al. 2022), broad-scale and coordinated conservation action will become increasingly more important and necessary to protect vulnerable species and habitats (Armsworth et al. 2015, Rhodes et al. 2022). Because of the need for conservation at a broad scale, global and national lists of at-risk species and conservation status frameworks have proliferated; however, they are only useful at the scale they are developed (Mace et al. 2008; Garnett et al. 2020; Naujokaitis-Lewis et al. 2022; Braulik et al. 2023). Moreover, these data sources may only focus on the rarest or most endangered species (Hamilton et al. 2024) leaving gaps in understanding of other declining species that may be of regional concern.

Since species are not limited by political borders, many have recommended regional-level landscape conservation and coordination as an effective way to tackle conservation planning (Lewis et al. 1996, Aldridge et al. 2008, Boyd et al. 2008), instead of focusing on more restricted and localized actions (AFWA 2021). Moreover, monitoring the response of species to threats and conservation interventions is most effective at the regional biome level (Bevan et al., 2024). However, regional conservation requires localized data, agreement on species taxonomies, and dedicated funding to develop and implement conservation strategies at the appropriate regional scale (Hamilton et al. 2024). Funding for conserving species also needs to address threats at a sufficiently large scope to be both successful and cost-effective, while also prioritizing species and actions that maximize investment return (Martin et al. 2018). However, broad conservation designs must be implemented on a much more local scale in specific locations, requiring discrete and iterative planning and implementation (Pressey et al. 2013).

While funding and legal mechanisms vary worldwide, in the United States, states have primary authority and responsibility for protecting and managing fish, wildlife, and plants (and their habitats) while also driving effective conservation strategies at local scales (USFWS et al. 2016). The Public Trust Doctrine, a legal doctrine that outlines the protection of public resources, reinforces this jurisdictional purview, emphasizing the role of each state in safeguarding natural resources to benefit current and future generations (Batcheller et al. 2010, Organ et al. 2012). In the past 20 years, State Wildlife Action Plans (SWAPs) have provided foundational local data to inform species management and policy for proactive regional conservation action (Lauber et al. 2009, USFWS 2020). These plans have allowed for strategic species prioritization, ensuring that conservation initiatives are effective and evidence-based (Hamilton et al. 2024). Further, the SWAPs are the foundation of a funding stream to states to implement SGCN conservation using the State Wildlife Grants program administered by the USFWS. In the northeastern U.S., SWAPs address ecological challenges, such as habitat loss and climate change impacts on biodiversity (Lacher and Wilkerson, 2013; Mahoney et al., 2015; Lauber et al., 2009).

To address the need for regional coordination, localized data, and funding dedicated to regional-level conservation actions, 14 Northeast U.S. states’ (Connecticut, Delaware, Maine, Maryland, Massachusetts, New Hampshire, New Jersey, New York, Pennsylvania, Rhode Island, Vermont, Virginia, West Virginia, and the District of Columbia) fish and wildlife agencies have collaborated to highlight species for coordinated conservation action over the past ∼50 years. Since 1990, the Northeast Fish and Wildlife Diversity Technical Committee (NEFWDTC; hereafter, “Tech Committee”) of the Northeast Association of Fish & Wildlife Agencies, Inc. (NEAFWA) has coordinated and worked with these 14 agencies to create and periodically update a non-regulatory list, now referred to as the Northeast Regional Species of Greatest Conservation Need (RSGCN). The RSGCN list is built from state lists of SGCN to maintain a non-regulatory list of species to provide focus, resources, and collaboration for conservation of mutual species of conservation concern (and their habitats) for current and future generations in the Northeast (Hamilton et al. 2024).

The RSGCN list spotlights species with population declines or emerging issues for collective conservation action and enhances knowledge of a species’ range-wide distribution, imperilment status, threats, and needed actions. Moreover, regional conservation programs, such as the Northeast Regional Conservation Needs (RCN) Grant Program, have emerged as key tools in this effort, with peer-reviewed studies, regionally consistent survey and monitoring protocols and Best Management Practices (BMPS), and conservation plans implemented across the region by many partners, validating the success of these collaborative strategies (e.g., Roy et al. 2002, Anderson et al. 2023, Heilferty et al. 2023, Skorupa et al. 2024). The impact of the RSGCN collaborative prioritization, particularly in the Northeast, has been significant, driving landscape-level conservation efforts and augmenting the implementation of SWAPs since 2005 at the regional scale. Due to the success of the Northeast RSGCN process, both the Southeast and Midwest U.S. regions have recently developed RSGCN lists (Rice et al. 2021, Radcliffe 2023). Based on the benefits of this process and subsequent adaptation in other regions, here, we present the methods, including recently updated methods from the compilation of the 2023 RSGCN list, with the following objectives: 1) document the RSGCN process, 2) summarize the benefits and outcomes of the process, and most importantly, 3) provide the blueprint for others to use across other regions.

## Methods

### Geographical Extent

The Tech Committee created the RSGCN list and dataset to provide non-regulatory data and related tools for regional species that states share and to prioritize and focus conservation in northeastern North America. For regional conservation, this included the jurisdictions of the 13 Northeast United States (Connecticut, Delaware, Maine, Maryland, Massachusetts, New Hampshire, New Jersey, New York, Pennsylvania, Rhode Island, Vermont, Virginia, and West Virginia), the District of Columbia, and six Canadian provinces (New Brunswick, Newfoundland and Labrador, Nova Scotia, Prince Edward Island, Quebec, and Ontario) within NEAFWA. The region includes the four Great Lakes that border at least one NEAFWA jurisdiction (i.e., Lakes Superior, Huron, Erie, and Ontario) and the Exclusive Economic Zone for the NEAFWA region, a 200 nautical mile buffer into marine waters (see Figure 1).

**Figure 1.**
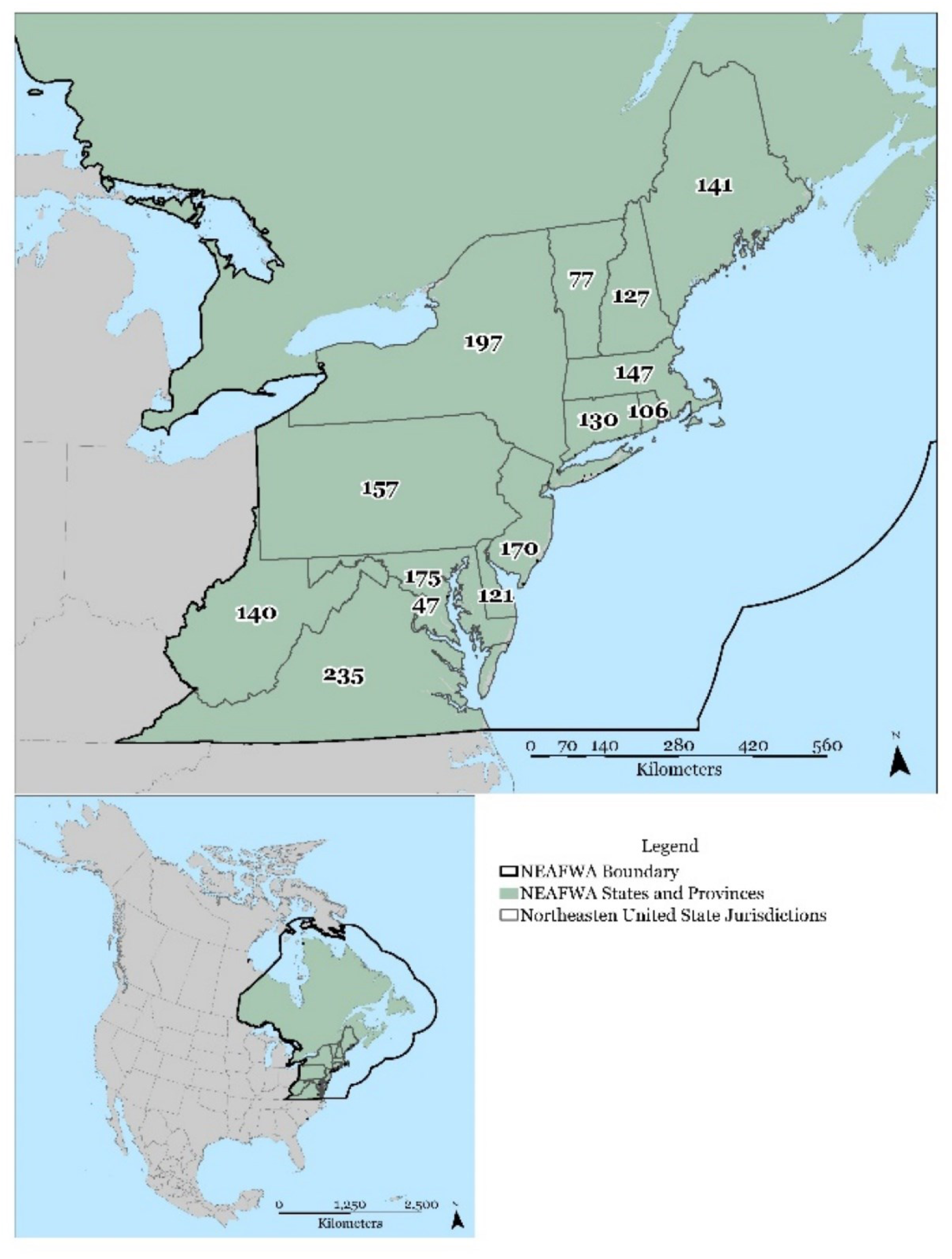
Map of the 14 Northeast jurisdictions with the number of RSGCN in each. Inset map below shows the boundary area of NEAFWA across North America.

The Northeast U.S. sustains more than 17,000 plant and animal species (NatureServe, 2021) and is home to some of the most densely populated states in the country. The terrestrial landscape of the Northeast U.S. is over 60% forested, with an average forest age of 60 years, and the region contains more than 200,000 miles of rivers and streams, nearly 37,000 water bodies, and more than 11.6 million acres of wetlands (Anderson et al. 2023, Olivero-Sheldon and Anderson 2008). Eleven globally unique habitats, from sandy barrens to limestone glades, support 2,700 restricted rare species (Anderson and Olivero-Sheldon 2011).

### RSGCN List

For the 2023 RSGCN list update, we implemented a 3-phase strategy (detailed below), meeting with members of the Tech Committee, their contractors, the 14 northeast SWAP coordinators, the Invertebrate Overview Team, and the RSGCN method group, who all provided the structure and guidance to solicit technical input from 20 teams of taxonomic experts (hereafter, “Taxa Teams”; see Appendix S3 for a list of team members for 2023). We conducted all analyses and data extraction (where noted) in R version 4.1.2 (R Core Team 2021) in RStudio (R Studio Team 2020).

#### Phase 1 – Data “Pre-population” Stage

Phase 1 of the RSGCN process involved data collection from numerous sources to generate a comprehensive list of all the wildlife species known to occur within the NEAFWA region (n = 17,923). Hereafter, the word “species” includes species, subspecies, or unique populations of management interest. To create this comprehensive list, we used the R package natserv (Chamberlain, 2020) to generate a list of all the species occurring within the study area. We extracted Species of Greatest Conservation Need (SGCN) lists from each SWAP (Appendix S4), previous versions of the Northeast RSGCN list, and RSGCN lists from the Midwest and Southeast regions (Terwilliger et al. 2015, Terwilliger et al. 2021, Radcliffe et al. 2023). We used the Environmental Conservation Online System (http://ecos.fws.gov/ecp/, Nov. 2021) to extract a list of all federally listed species that occur in the NEAFWA region and any species on the National Listing Workplan that occur in the NEAFWA region (USFWS 2021). We collected lists of state-protected species from the websites of state wildlife agencies, heritage programs, marine programs, and other natural resource departments, depending on the state and taxonomic group, state code, and statutes (See Appendix S4). We also incorporated a recently updated dataset for Plecoptera, Trichoptera, and Ephemeroptera (Terwilliger et al. 2021) to expand the list of included species from these orders since these data were more comprehensive than those available on NatureServe.

We used multiple spatial datasets to identify additional species for inclusion in our comprehensive list. We used NatureServe Explorer, the International Union for the Conservation of Nature (IUCN) Red List of Threatened Species, the Ocean Biodiversity Information System (OBIS), and the Botanical Information and Ecology Network (BIEN) (NatureServe 2021, IUCN 2021, OBIS 2021, BIEN 2021). Other publicly available spatial data (e.g., iNaturalist and Global Biodiversity Information Facility) are available, but we opted not to use these sources because they are point data rather than complete distributions. OBIS data are point data extrapolated to grid cells, but it was the best available source we found for marine species, which are largely absent from the other sources we used. For the IUCN Redlist data, we downloaded spatial data files for all relevant taxonomic groups from their Spatial Data Download page (IUCN 2021). For OBIS and BIEN, we used the R packages robis (Provoost and Bosch 2017) and BIEN (Maitner et al. 2022) to extract spatial data. For all these data layers, we extracted any species whose distribution layer overlapped with the study area (Figure 1).

We combined the list of species names from all sources listed above and removed duplicates. To address taxonomic synonymies between the lists, we used the R package taxize (Chamberlain et al. 2020) to match each scientific name to its associated Taxonomic Serial Number (TSN) from the Integrated Taxonomic Information System (ITIS; ITIS 2021, Blum & Chamberlain 2024). We extracted the TSN, scientific name, and any known synonyms for each species from ITIS. All species in our comprehensive list who shared an accepted TSN and name were synonymized into a single record unless: (a) two species that are considered synonymous in ITIS were both included in a single state’s protected species or SGCN list, (b) an invalid taxonomic entity is currently listed at the federal level or is being considered for listing, or (c) the accepted name in ITIS has been superseded by a taxonomic authority or a recent publication (e.g. Crandall and DeGrave 2017). For any species that did not have a matching species-level record in ITIS, we manually generated a TSN comprised of the TSN for the nearest higher taxonomic entity and a letter assigned sequentially. For example, the mayfly *Ameletus janetae* has no record in ITIS; the TSN for the genus *Ameletus* is 100996, so the generated TSN for *Ameletus janetae* is 100996a. This ensured every species in our comprehensive list had a unique TSN ID number that could be used throughout the RSGCN process.

Once we reviewed the comprehensive list, we excluded species that only occur in the Canadian portion of NEAFWA, are extirpated from the region, are extinct, or are non-native. We extracted taxonomic information using R package ritis (Blum & Chamberlain 2024) and both conservation status (Appendix S4) and state-level occurrence using R package natserv (Chamberlain 2020). We manually entered taxonomic information for species with generated TSN IDs. We considered a species to occur in a state if it had any Subnational Rank (hereafter, “S-rank;” a metric used by NatureServe to assess extirpation risk – see Master et al. 2012) except those listed as possibly extirpated or presumed extirpated in NatureServe, was included on a state’s regulatory list, or was included in a SWAP as an SGCN. For Plecoptera, Trichoptera, and Ephemeroptera species, we used data Terwilliger et al. (2021) from whenever it superseded NatureServe occurrence information. We calculated an average S-rank value for each species based on its S-rank in all northeast states. We also summed the number of states where each species is on a regulatory or SGCN list.

Using these data, we pre-populated draft taxon lists for each Taxa Team to review. We applied selection criteria to produce species lists into four categories: Likely RSGCN, Maybe RSGCN, Not Likely RSGCN, and Unknown RSGCN. This prescreening effort helped to organize and prepare the data for more efficient review by taxa experts. First, we filtered each species by its Global Rank (i.e., G1 and G2) and further filtered the data through their U.S. federal listing status (e.g., endangered, candidate species). Next, we filtered each species by their average northeast regional state S-rank, with a cut-off of an S-rank of less than S2. Lastly, we filtered each species by State Protected Status (i.e., state endangered or threatened). It is important to note that we may have identified a species as either Likely or Maybe SGCN if it didn’t meet one of the above criteria but met others.

We initially assigned species to a regional Concern Level based on its conservation status at global, national, and state scales (see Appendix S4 for details about these ranking systems). We assessed each species and assigned them a Regional Concern Level based on a relative measure of species imperilment of Very High, High, or Moderate Regional Concern Level based on their conservation status.

We then calculated the Regional Responsibility (the proportion of a species’ range that falls within the boundaries of the NEAFWA region) for each species using the spatial data extracted from NatureServe Explorer, IUCN Red List, OBIS, and BIEN. We determined the Regional Responsibility by dividing the area of the species range within the study area by the total area of the study area to get the computed percent of regional responsibility. If a species had distribution layers available from multiple sources, we assigned the regional responsibility based on the maximum value. For species with no spatial distribution data, we calculated a proxy for regional responsibility by dividing the number of NEAFWA states and provinces where the species occurs by the total number of states and provinces in which it occurs. Out of the 806 species, 764 had enough data to predict Regional Responsibility from the sources listed above before we met with the Taxa Teams. We assigned each species to one of the Regional Responsibility categories (<25%, 25-50%, 50-75% 75-100%, or 100% [endemic]). Any species that did not have distribution data or state occurrence information was assigned “Unknown” for the Regional Responsibility.

Lastly, we added fields to the dataset, such as taxon and subtaxon fields based on species taxonomy, life history traits that might be a limiting factor for the species, habitat use information, and threats (see Terwilliger et al. 2023 for details), and used these data fields to create taxon-specific datasets for review of each of the Taxa Teams.

#### Phase 2 – Incorporating Regional Expertise Data Stage

The RSGCN methodology subcommittee of the NEFWDTC and an advisory Invertebrate Overview Team selected 20 taxonomic groups for which sufficient regional expertise exists to evaluate for the 2023 RSGCN list. To incorporate input from regional experts from all Northeast states into the RSGCN process, we compiled worksheets (see Appendix S5 for an example) for each of the 20 Taxa Teams, representing approximately 175 individuals, which included state biologists (n=140) and external experts from partner organizations and academia (n=35), which came “pre-populated” with the data we collected in Phase 1. Taxa Team members reviewed the worksheets, indicating whether they agreed with the “pre-populated” values for concern level, regional responsibility, and RSGCN status. They also provided suggestions for overriding factors, notes on species taxonomy, and other comments explaining their decisions. Once we received the completed worksheets from each Taxa Team member, we compiled their results into a single document. Each of the 20 Taxa Teams then met to discuss the initial rounds of reviews and vote on which candidate species should be considered as an RSGCN or assigned as a “Watchlist” species, which are species that either do not have sufficient data to decide, have varying population trends in different parts of their range, is interdependent with an existing RSGCN but do not meet the requirements on its own, or has little Regional Responsibility but is of particularly high concern. During this process, Taxa Teams identified 63 species missed by the prescreening process that they listed in one of the six categories (e.g., RSGCN, Watchlist). Some teams met more than once (n = 4), and some (n = 2) contributed additional input to their documents after their meetings. All data, comments, and votes were compiled and incorporated into the RSGCN List and database as the new draft RSGCN list.

#### Phase 3 – Approval Process Stage

The last phase of the RSGCN process required the review and approval of all the Taxa Teams, the Tech Committee, other NEAFWA technical committees, and a series of NEAFWA committees and state agency directors before finalizing the list (Figure 2). Once the Taxa Teams reviewed and approved each taxon’s RSGCN list at the end of Phase 2, we sent the list to the NEFWDTC for preliminary review. Once the NEFWDTC provisionally approved the RSGCN list, we sent the RSGCN List to other NEAFWA technical committees (Upland Gamebird Technical Committee, Fur Resources Technical Committee, and Rivers and Streams Technical Committee) for review, and they added more data to four species records and recommended the status change of three species. These changes were reviewed by the Taxa Teams and NEFWDTC, which approved the status change of two of the three species identified by the other committees. We then sent them to NEAFWA’s Administrators and Directors. Once approved by the Directors, the RSGCN list became final, with two species changing statuses through the review and approval process.

**Figure 2.**
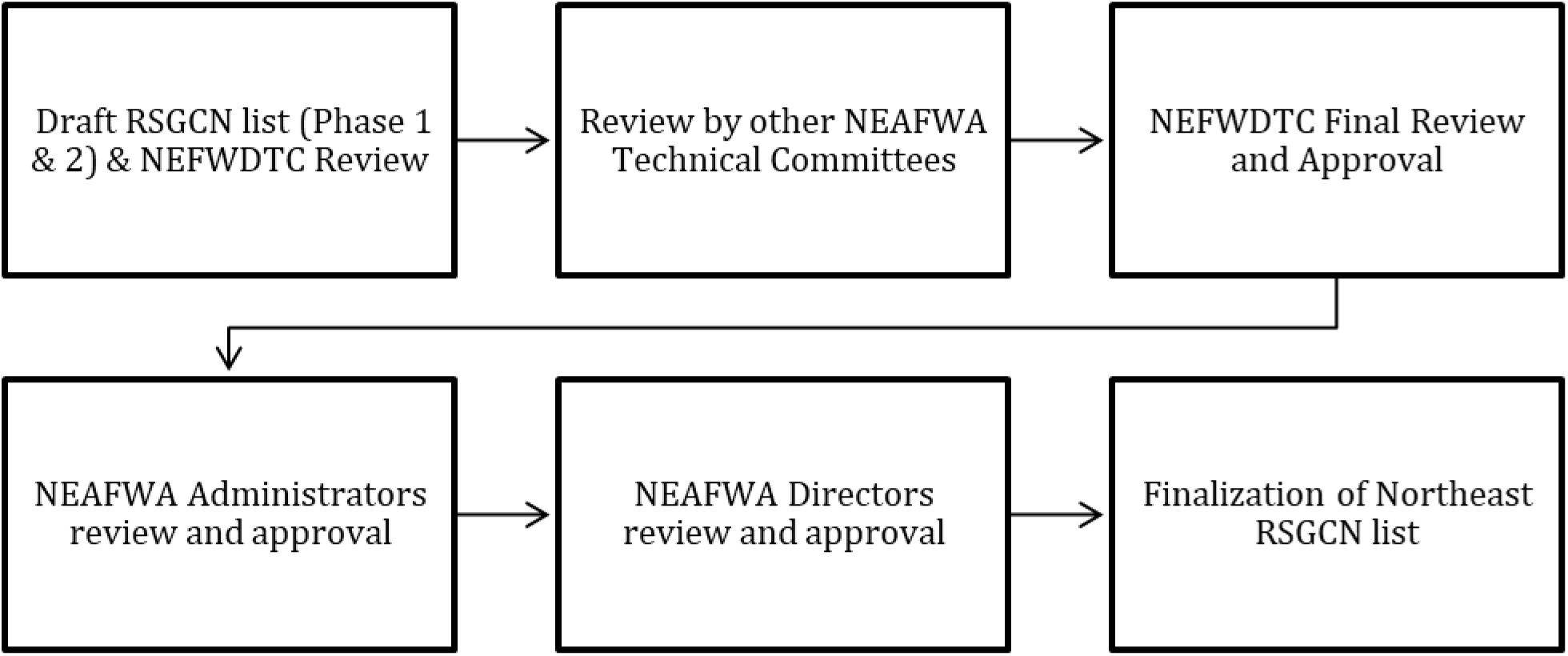
Flow chart of approval process.

### Analyses

#### 2023 RSGCN List Summary

We summarized the taxonomic group breakdown and RSGCN categories across the region by comparing the total percent of species in the region to those on the RSGCN list and then comparing taxon numbers in each category.

#### How well did the global, national, and state assessments predict RSGCN status?

Since pre-populated data came from global, national, and state-level datasets, and the goal of RSGCN is for regional conservation, we tested how well these datasets compare to the regional data after more localized Taxa Team input to evaluate their efficacy. To assess the relative effectiveness of the pre-population process (Phase 1), using the tidyverse package (Wickham et al. 2019), we compared the initial predictions of RSGCN status, Concern Level, and Regional Responsibility to the final decisions of the Taxa Team experts (Phase 2) by taking the predicted value for each category and calculating the percentage of predictions that matched the final decisions for RSGCN status, Concern Level, and Regional Responsibility.

## Results

### 2023 RSGCN List Summary

Out of the 17,923 Northeast species, we evaluated and prescreened 7,270 using the NEAFWA RSGCN selection criteria within the 20 taxonomic groups. Of the 7,270, we identified 1,817 as SGCN (Species of Greatest Conservation Need) in Northeastern U.S. 2015 SWAPs. Of these SGCN, we identified 693 invertebrates and 230 were plants beyond the scope of this assessment due to lack of jurisdiction and expertise across the entire taxon. However, we flagged these species for inclusion in the next 2028 RSGCN update. The analysis resulted in 382 RSGCN, 36 Proposed RSGCN (Appendix S1), 229 Watchlist Assessment Priority, and 61 Proposed Watchlist Assessment Priority (Appendix S1), for a total of 806 species. Of all Northeast species initially considered for the RSGCN list, only 11% met the RSGCN criteria and were assigned to one of the RSGCN list categories (Table 4). Of the total species, 806 species met at least one of the RSGCN status categories and 47.2% (n= 382) of those meeting the criteria for RSGCN status (Figure 3). The two “Proposed” categories do not qualify as RSGCN since they are not currently listed as SGCN in any Northeast SWAP and represent 12.3% (n = 97) of the 806 species total. The RSGCN Watchlist [Assessment Priority] category (new to the 2023 method revision, Appendix S2) contains 28.2% (n = 228) of listed species highlighting species with data deficiencies, taxonomic uncertainties, or variable trends within the region. Three species met RSGCN Watchlist [Interdependent Species] criteria, and 95 additional species were deferred to other regions for interregional coordination and primary stewardship in the core of their range.

**Figure 3.**
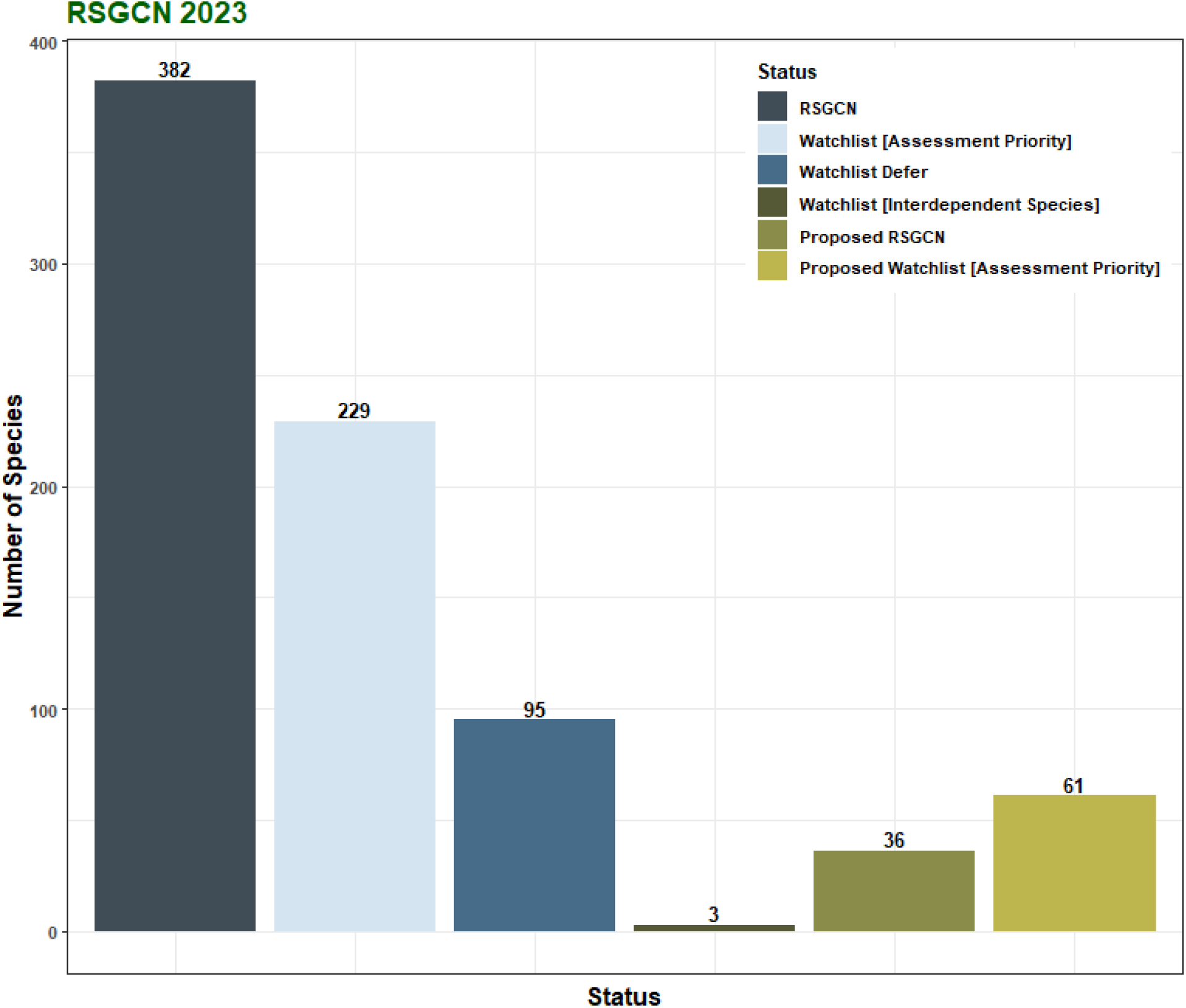
Number of RSGCN and Watchlist species in each category.

### RSGCN & Watch List Species Lists

National and global datasets captured 38.5% (n = 310) and 38.8% (n = 313) of Final RSGCN and Watchlist species (n = 806) as either Likely or Maybe RSGCN, respectively, flagging Taxa Teams to assess the species for the Regional List (Table 1, Figure 4). From the total number of northeast species assessed (n = 17,923), 33.9% (n = 6077) of those species were initially identified as Likely and Maybe categories and were not listed as RSGCN or Watchlist species by regional state experts. RSGCN were predicted correctly 54.5% (n = 208) of the time, whereas expert opinion confirmed 33.8% (n = 129) of species in the Maybe category from pre-screening and added 11.8% (n = 45) of species from either the Not Likely RSGCN category or species that were not pre-screened from their discussions of current data and trends across the region.

**Figure 4.**
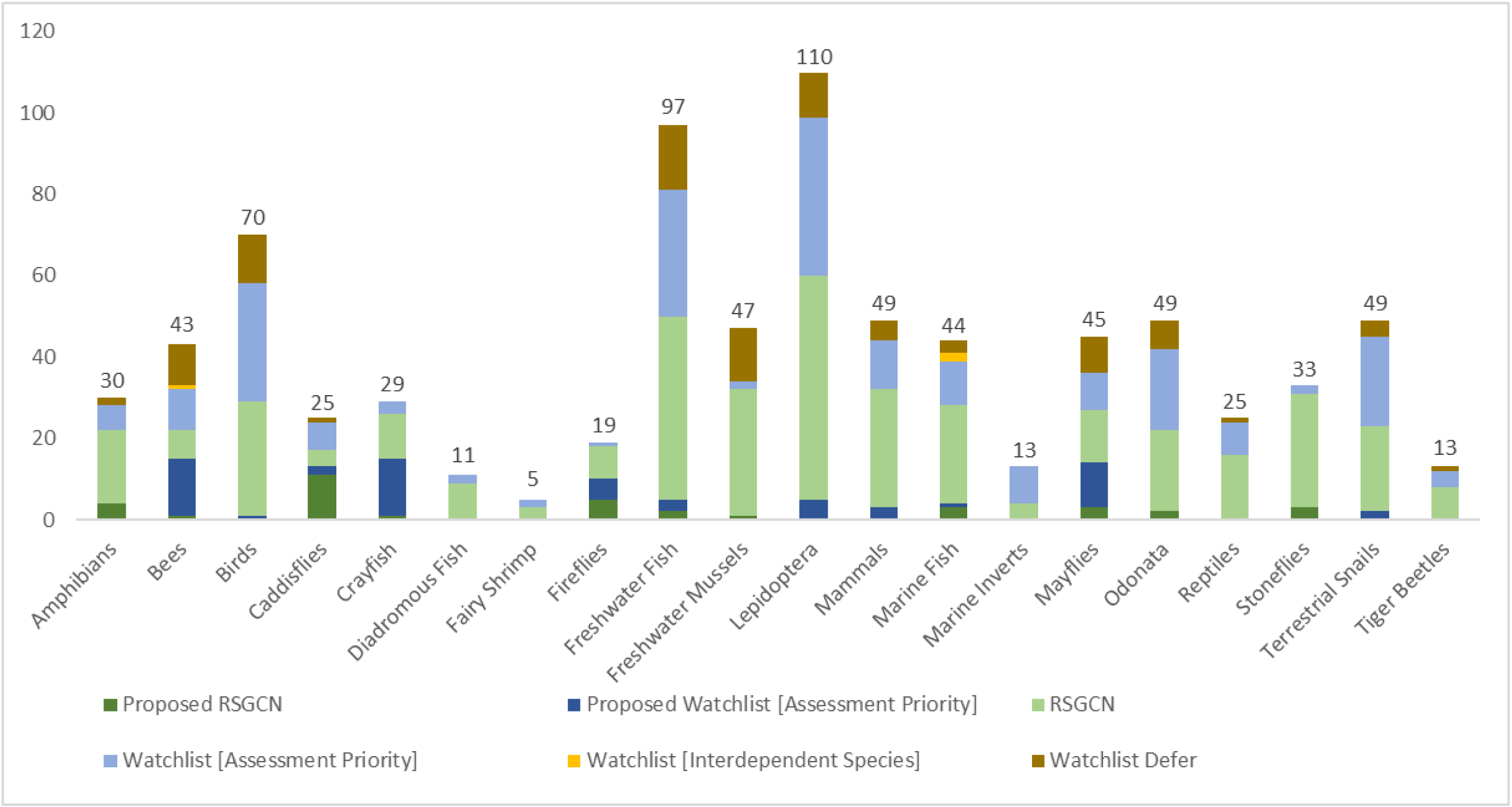
The 20 taxonomic groups that Taxa Teams assessed, along with the total number of each taxon across the top, and the RSGCN & Watchlist categories are shown in color.

**Table 1.** Rows show the predicted values from the pre-population process compared to the final status of regional species in greatest conservation need. For example, out of the 382 final NE RSGCN species, 54.5% of them were predicted to be Likely RSGCN from the pre-population process.

|  | Not RSGCN |  | Proposed RSGCN |  | Proposed Watchlist [Assessment Priority] |  | RSGCN |  | Watchlist [Assessment Priority] |  | Watchlist [Interdependent Species] |  | Watchlist Defer |  | Total |
| --- | --- | --- | --- | --- | --- | --- | --- | --- | --- | --- | --- | --- | --- | --- | --- |
| Likely RSGCN | 1473 | 8.6% | 20 | 55.6% | 9 | 14.8% | 208 | **54.5%** | 63 | 27.5% | 0 | 0.0% | 10 | 10.5% | 1783 |
| Maybe RSGCN | 4604 | 26.9% | 6 | 16.7% | 10 | 16.4% | 129 | 33.8% | 95 | 41.5% | 0 | 0.0% | 73 | 76.8% | 4917 |
| Not Likely RSGCN | 10676 | 62.4% | 6 | 16.7% | 13 | 21.3% | 30 | 7.9% | 60 | 26.2% | 2 | 66.7% | 9 | 9.5% | 10796 |
| Not Pre-screened | 364 | 2.1% | 4 | 11.1% | 29 | 47.5% | 15 | 3.9% | 11 | 4.8% | 1 | 33.3% | 3 | 3.2% | 427 |
| **Final** | **17117** |  | **36** |  | **61** |  | **382** |  | **229** |  | **3** |  | **95** |  | **17923** |

### Regional Responsibility

Overall, the national and global datasets accurately predicted the Regional Responsibility of 78.2% (n = 598) of species with sufficient information for determination and analysis (n = 764). Taxa Teams corrected or added additional Regional Responsibility information for 25.2% (n = 202) of the final 806 species. Species with less than 25% of their range within NEAFWA were the most accurately predicted (80.3%), followed by endemic species (75.7%), with Taxa Team input correcting 19.7 to 30.1% of the other predicted values (Table 2). Six of the watchlist assessment priority species still had undefined ranges; research on their distribution is one of the reasons to list them as assessment priority species.

**Table 2.** Regional Responsibility Predicted Results before and after Taxa Team input (percent accuracy)

|  |  | Final |  |  |  |  |  |  |  |  |  |
| --- | --- | --- | --- | --- | --- | --- | --- | --- | --- | --- | --- |
|  |  | 100%<br>(NEAFWA<br>Endemic) |  | 75-100% |  | 50-75% |  | 25-50% |  | <25% |  |
| Predicted | 100%<br>(NEAFWA<br>Endemic) | **128** | **75.7%** | 10 | 10.3% | 8 | 5.0% | 2 | 1.1% | 2 | 1.0% |
|  | 75-100% | 14 | 8.3% | **67** | **69.1%** | 3 | 1.9% | 4 | 2.3% | 0 | 0.0% |
|  | 50-75% | 5 | 3.0% | 13 | 13.4% | **120** | **74.5%** | 19 | 10.9% | 7 | 3.5% |
|  | 25-50% | 5 | 3.0% | 4 | 4.1% | 20 | 12.4% | **124** | **70.9%** | 21 | 10.6% |
|  | <25% | 0 | 0.0% | 1 | 1.0% | 3 | 1.9% | 23 | 13.1% | **159** | **80.3%** |
|  | Total | 153 |  | 96 |  | 154 |  | 172 |  | 189 |  |
|  | **Final\*** | **169** |  | **97** |  | **161** |  | **175** |  | **198** |  |

\*Final numbers include species on the list not prescreened

### Regional Concern Level

The national and global datasets accurately predicted the Regional Concern Level for 57% (n=240) of species (Table 3). Taxa Teams input led to a 67% (n=52) increase in Very High-concern species, 73% (n=75) of High-concern species, and 86% (n=51) of Moderate-concern species missed by national and global datasets across 20 taxonomic groups (Figure 5).

**Figure 5.**
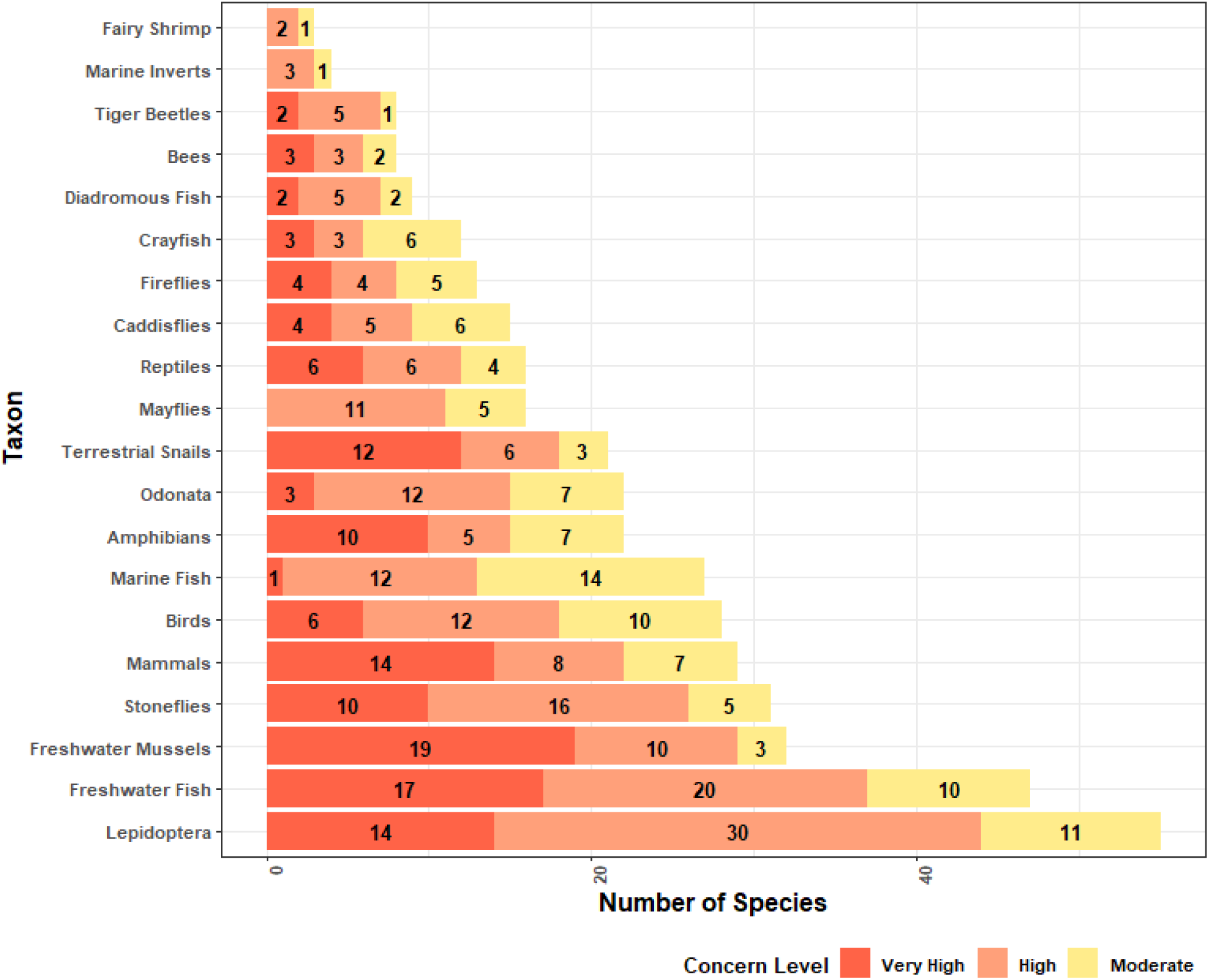
RSGCN and Proposed RSGCN regional concern levels are shown within each of the 20 taxonomic groups assessed.

**Table 3.** Regional Concern Level results for RSGCN and Proposed RSGCN before and after Taxa Team input (percent accuracy)

|  |  | Final |  |  |  |  |  |
| --- | --- | --- | --- | --- | --- | --- | --- |
|  |  | Very High |  | High |  | Moderate |  |
| Predicted | Very High | **78** | **60.0%** | 28 | 15.7% | 1 | 0.9% |
|  | High | 38 | 29.2% | **103** | **57.9%** | 22 | 20.0% |
|  | Moderate | 2 | 1.5% | 26 | 14.6% | **59** | **53.6%** |
|  | Low | 9 | 6.9% | 20 | 11.2% | 26 | 23.6% |
|  | Total | 127 |  | 177 |  | 108 |  |
|  | **Final \*** | **130** |  | **178** |  | **110** |  |

\*Final numbers include RSGCN & Proposed RSGCN not prescreened

**Table 4.** Sub taxon breakdown of species known to occur in the study region, number that are SGCN from the 2015 State wildlife Action Plans, number of species that are RSGCN and Watchlist species in the northeast region.

|  | **Northeast**<br>**Species** | **SGCN** | **RSGCN**<br>**(incl.**<br>**Proposed)** | **WL**<br>**Assessment**<br>**Priority**<br>**(incl.**<br>**Proposed)** | **WL**<br>**Defer** | **WL**<br>**Interdependent** | **All**<br>**RSGCN/WL**<br>**Categories** |
| --- | --- | --- | --- | --- | --- | --- | --- |
| *Birds* | 426 | 284 | 28 | 30 | 12 | 0 | 70 |
| *Mammals* | 183 | 107 | 29 | 15 | 5 | 0 | 49 |
| *Amphibians* | 111 | 88 | 22 | 6 | 2 | 0 | 30 |
| *Reptiles* | 115 | 84 | 16 | 8 | 1 | 0 | 25 |
| *Fish – Fresh* | 336 | 213 | 47 | 34 | 16 | 0 | 97 |
| *Fish –*<br>*Diadromous* | 28 | 14 | 9 | 2 | 0 | 0 | 11 |
| *Fish –*<br>*Marine* | 661 | 102 | 27 | 12 | 3 | 2 | 44 |
| *Terrestrial*<br>*Snails* | 268 | 182 | 32 | 24 | 4 | 0 | 60 |
| *Freshwater*<br>*Bivalves* | 149 | 106 | 21 | 2 | 13 | 0 | 36 |
| *Crayfish* | 78 | 26 | 12 | 17 | 0 | 0 | 29 |
| *Fairy, Clam,*<br>*& Tadpole*<br>*Shrimp* | 17 | 5 | 3 | 2 | 0 | 0 | 5 |
| *Dragonflies*<br>*and*<br>*Damselflies* | 255 | 205 | 22 | 20 | 7 | 0 | 49 |
| *Butterflies*<br>*and Skippers* | 224 | 134 | 26 | 12 | 5 | 0 | 43 |
| *Moths* | 2426 | 364 | 29 | 32 | 6 | 0 | 67 |
| *Tiger Beetles* | 40 | 35 | 8 | 4 | 1 | 0 | 13 |
| *Fireflies* | 44 | 13 | 13 | 6 | 0 | 0 | 19 |
| *Caddisflies* | 565 | 40 | 15 | 9 | 1 | 0 | 25 |
| *Mayflies* | 281 | 62 | 16 | 20 | 9 | 0 | 45 |
| *Stoneflies* | 253 | 67 | 31 | 2 | 0 | 0 | 33 |
| *Bumble Bees* | 23 | 17 | 3 | 3 | 4 | 0 | 10 |
| *Solitary Bees* | 400 | 131 | 5 | 21 | 6 | 1 | 33 |
| *Marine*<br>*Invertebrates* | 466 | 95 | 4 | 9 | 0 | 0 | 13 |
| *Plants* | 6084 | 1785 | n/a | n/a | n/a | n/a | n/a |
| *Other*<br>*species* | 4490 | 632 | n/a | n/a | n/a | n/a | n/a |
| *Total* | 17923 | 4788 | 418 | 290 | 95 | 3 | 806 |

## Discussion

Overall, the national and global datasets were only able to accurately identify 55% of species found on the final RSGCN list, 78% of the Regional Responsibility, and 57% of the Regional Concern Level. Without the local expertise of the 20 Taxa Teams sharing data from each state across the region, merely relying on these databases to assess regional prioritization would have resulted in hundreds of species being overlooked and much poorer data quality overall. Since access to the best available data is the foundation of sound conservation prioritization and decision-making, and coordination among various organizations is essential moving forward, especially in the face of mounting uncertainty from climate change (Armsworth et al. 2015), the benefits of regional collaborative sharing of current data this process cannot be overstated. Thus far, two other U.S. regions, the Midwest and Southeast, have adopted the RSGCN process for their own regional conservation planning and implementation needs (Rice et al. 2019, Terwilliger et al. 2021, Radcliffe et al. 2023).

While national and global species lists, such as the IUCN Red List, are extremely important for conservation, non-regulatory regional lists (like RSGCN) fill the much-needed landscape conservation gap to help proactively designate and collaborate on species and habitat conservation at the regional scale before species are federally listed. One key benefit of the RSGCN process is identifying (and prioritizing) species before they are candidates for regulatory protection since conservation projects to prevent extirpation or extinction are generally more cost-efficient and effective with more conservation partners and landowners willing to work to preclude listing. For instance, regional collaborative conservation efforts in the Northeast for New England Cottontail (*Sylvilagus transitionalis*), Brook Floater (*Alasmidonta varicosa*), Lake Sturgeon (*Acipenser fulvescens*), and Delmarva Fox Squirrel (*Sciurus niger cinereus*) are all contributing factors cited by the USFWS in determinations to not list or to delist each species under the Endangered Species Act (USFWS 2015a, 2015b, 2018 & 2024, Fuller and Tur 2012).

Another important step in implementing regional- and landscape-level conservation is setting up funding mechanisms for multi-state projects (Hamilton et al. 2024). Since 2007, the Tech Committee has allocated over $7.7 million through the RCN Grant Program to landscape-scale conservation projects focused on these RSGCN-identified priorities in the Northeast, leveraging an additional $6.6 million in conservation investment from partners across at least 114 RCN Grant projects on 39 Competitive State Wildlife Grant (CSWG) projects from 2008 to 2023, with $19.1 million in awards amplified by $8.25 million in matching funds for a total investment of $27.36 million in regional conservation efforts. The minimum total regional conservation collaborative investment exceeds $41.6 million since the RCN program began in 2007, serving as seed funds for RSGCN-related research and conservation implementation by various federal agency partners and academia (e.g., U.S. Fish and Wildlife Service, U.S. Geological Survey, U.S. Department of Defense, Cornell University, and University of Massachusetts Amherst).

An example of one of these RCN grants was the award of $72,940 in 2012 which initiated and funded a range-wide status assessment of the Brook Floater (Wicklow 2017). The report spurred the creation of a Brook Floater Working Group, now comprised of 39 representatives from 3 federal and 14 state agencies and four academic institutions. Since the initial RCN grant, the working group won multiple Competitive State Wildlife Grants (∼$1.5 million in total), resulting in rapid and long-term assessment protocols (Sterrett et al. 2018, Sterrett et al. 2022), peer-reviewed publications (e.g., Skorupa et al. 2024), a structured decision-making workshop to identify locations for reintroduction (Roy et al. 2022), and propagation for species recovery across its range. A similar ongoing series of regional conservation efforts for the Chesapeake Logperch (*Percina bimaculata*) are in the process of determining its extant distribution, characterizing genetic diversity, developing a conservation action plan, initiating a captive rearing program, actively augmenting extant populations, and reintroducing the fish in its historic range (for the initial status review for the Chesapeake Logperch, see NEFWDTC [2018]).

Other programs kickstarted by RCN Grants have had similar success, such as the combined $5.4 million in conservation funding for six RSGCN freshwater turtles over the past two decades from RCN and CSWG Grant Programs, which has since leveraged a total of nearly $10 million to support conservation actions for this group of RSGCN turtles and their habitats. These species niches are highly partitioned in the Northeast and the six RSGCN turtle species serve as flagship species for various priority habitat types, from calcareous fens to cold water streams (Compton 2007, Jones et al. 2018, Willey et al. 2022, Erb and Roberts 2023). This funding has established or supported multiple turtle-specific working groups (e.g., Northeast Partners in Amphibian and Reptile Conservation and Diamondback Terrapin Working Group). Species working groups have been established through the Northeast Partners in Amphibian and Reptile Conservation and Diamondback Terrapin Working Group with biologists representing state and federal fish and wildlife agencies, academia, and non-governmental organizations collaborating on multiple regional status assessments regionally consistent protocols, BMPS, and conservation plans for all six RSGCN turtles, many using a shared framework for freshwater turtle conservation planning in the region. To date, these working groups also have collaborated to create, enhance, and restore nesting habitat, headstart thousands of juvenile turtles, establish the Collaborative to Combat the Illegal Trade in Turtles and create genetic databases to assist in the geolocation and repatriation of confiscated turtles. An ongoing RCN-funded project is implementing high priority actions identified in the RCN-funded Northern Diamondback Terrapin Conservation Strategy (*Malaclemys terrapin terrapin*; Egger and the Diamondback Terrapin Working Group 2016) and Northeast SWAPs, coordinating a regional population status assessment using standardized survey techniques across seven coastal states, which will be used to create a shared regional data portal, identify and map a state and regional terrapin conservation area network, and inform the spatial ecology of the RSGCN turtle.

Northeast regional conservation projects that address the needs of RSGCN have focused on improving habitat conditions. A recent RCN-funded project aimed to improve management of regionally significant xeric grasslands, barrens, and woodlands habitat for pollinators in the Northeast by developing survey protocols and conducting habitat management on more than 740 acres at 20 sites across ten states (Heilferty et al. 2023). Associated monitoring for bees, moths, and vegetation communities to measure the effectiveness of different habitat treatments and develop best management practices collected approximately 81,000 records of 1,709 species of bees and moths (Heilferty et al. 2023). Another regional project coordinated the installation and repair of gates and temperature improvement structures on 11 caves and mines known to support RSGCN and SGCN bat hibernacula in four states (USFWS 2020). An early RCN-funded project to create an inventory of aquatic connectivity barriers in the Northeast (Martin and Apse 2011) has expanded since then to include information for at least 29,583 dams, which has informed the removal of at least 346 dams and restored more than 3,657 miles of aquatic connectivity in the last decade (Anderson et al. 2023).

While the RSGCN list has been important in generating and securing funding for regional conservation projects and fostering collaborations across states and sectors, one of the most important conceptual contributions of this RSGCN list is that it recognizes the role of the state fish and wildlife agencies and their collaboration while considering other criteria and ranking scales and systems. This results in a bottom-up, state-driven proactive species prioritization and conservation effort at a regional scale that informs planning at the state, regional, and national levels. At the state level, the RSGCN list is used to identify SGCN in SWAPs, informing the regional context for the plans and identifying lists of priority species in taxonomic groups for which a state fish and wildlife agency may have limited expertise or capacity (AFWA, 2021). Regionally, the development of Northeast RSGCN has led to the development of RSGCN in other regions (e.g., Rice et al. 2019, Terwilliger et al. 2021, and Radcliffe et al. 2023). The Northeast Region of the USFWS included RSGCN as a primary selection criterion in the designation of Northeast At-Risk Species (USFWS 2021). RSGCN species are used to guide various regional conservation plans, such as the U.S. Geological Survey’s syntheses of the effects of climate change on RSGCN and climate-smart conservation actions for RSGCN (Staudinger et al. 2015 & 2024) and State Forest Action Plans (United States Forest Service and Northeast-Midwest State Foresters Alliance 2022).

While the examples of the coordination and funding spurred on by the RSGCN process listed above help demonstrate the utility of this approach, we acknowledge that it is not a simple solution for developing the best available data for a region or fostering collaboration across municipalities and sectors. Bringing together hundreds of volunteer experts from multiple state agencies, universities, non-profits, and other organizations is a difficult task that requires extensive planning, coordination, and patience. The methods presented here result from hundreds of people working together to refine the process over many years, and while we do not doubt there is room for improvement, the benefits of the RSGCN list have been invaluable for both the states and the region, they have been especially vital for the vulnerable species of the Northeast.

## CONSERVATION IMPACT SUMMARY RESULTS

The iterative process of identifying species with the greatest conservation need for the northeast region of the United States using information gathered and verified by regional state experts led to greater cooperation among agencies and organizations across state borders. It catalyzed funding and concrete conservation actions to improve the knowledge and status of species and habitats, proving successful in both landscape- and species-level conservation.

## ACKNOWLEDGEMENTS

We acknowledge the contribution of the 200+ individuals who have contributed to RSGCN conservation in the Northeast as part of the NEFWDTC, SWAP Coordinators, and/or 2023 Taxa Teams from 2018 to 2023, on which this article is based (see northeastwildlifediversity.org/rsgcn or Appendix S3 for a full list). In addition, we also acknowledge the support of the NEAFWA Region State Directors for their help in reviewing and supporting this work and submission. Some information was provided by NatureServe (www.natureserve.org) and the NatureServe Network, a leading source of information about rare and endangered species, and threatened ecosystems.

## Supporting Information

Additional supporting information may be found in the online version of the article at the publisher’s website.

## APPENDIX S1. Pre-screening results: excel spreadsheet with the following data fields

1. Scientific Name
2. Common Name
3. Pre-screened RSGCN category
4. Final RSGCN category
5. Pre-screened Regional Responsibility
6. Final Regional Responsibility
7. Pre-screened Concern Level
8. Final Concern Level

## APPENDIX S2. Method definitions

## APPENDIX S3. Table of all contributions from Taxa Team members

## APPENDIX S4. Excel book (or word tables) of data sources and types

## APPENDIX S5. Example Taxonomic Team Worksheet

