## Appendix S2, S3, S4 for "Strategic Conservation Prioritization Through Creating and Updating the Regional Species of Greatest Conservation Need (RSGCN) List in the Northeastern U.S."

### Appendix S1

#### RSGCN Pre-screening results

Shared as an Excel workbook

1. Scientific Name
2. Common Name
3. Pre-screened RSGCN category
4. Final RSGCN category
5. Pre-screened Regional Responsibility
6. Final Regional Responsibility
7. Pre-screened Concern Level
8. Final Concern Level

### Appendix S2

#### Definitions for RSGCN Methods

- **RSGCN**: regional species of greatest conservation need, Native species for which the Northeast region has a stewardship responsibility due to high conservation concern and/or populations that are concentrated in the Northeast Region and that have been identified as SGCN by at least one Northeast state. (Terwilliger Consulting Inc. & Northeast Fish and Wildlife Diversity Technical Committee 2022).
- **Regional Responsibility:** the proportion of the species' North American or North Atlantic range overlapping the NEAFWA region (including the Canadian Provinces of Ontario, Quebec, New Brunswick, Nova Scotia, Newfoundland, Labrador, and Prince Edward Island), calculations were refined.
- **Concern Level:** indicates the level of conservation status and need in the region, which are Very High, High, and Moderate.
- The formalization of **Regional Responsibility Overriding Factors (ROF)** and **Concern Overriding Factors (COF)**. ROF and COF were identified by the taxa teams to document the reasons for identifying a species as RSGCN to clarify RSGCN status when it does not otherwise meet either the Regional Responsibility or Concern selection criteria.
- **Regional Responsibility Overriding Factors** include:
    - ***Core Population***: Species found over a very large geographic area, but the strongest populations are in the NEAFWA region.
    - ***Climate Change Range Shift***: Species where predicted range shifts due to climate change would make the species a higher regional responsibility in the future.
    - ***Migratory Species***: Species where the overall geographic range does not meet the 50% threshold for regional responsibility, but specific seasonal ranges do. Migratory species may be included as RSGCN if:
        - $\geq$ 50% of the breeding range occurs in the Northeast (the NEAFWA region, including Canadian Provinces)
        - $\geq$ 50% of the migratory stopover habitat occurs in the Northeast
        - $\geq$ 50% of the wintering habitat occurs in the Northeast
    - ***Highly Imperiled***: Species is highly imperiled throughout its range and is of high conservation concern in every region in which it occurs.
    - ***Disjunct Population:*** Species has a disjunct population that may contribute to genetic diversity or the three R's (resiliency, redundancy, or representation) when conducting species status assessments.
    - ***Stewardship Priority:*** The region has a significant stewardship responsibility for management, restoration, or recovery of the species.
- **Concern Overriding Factors** include:

    - ***Emerging:*** Species where conservation statuses are likely to change quickly as a result of a new or widespread threat, such as disease or a shift in market forces driving harvest or collection.
    - ***Climate Vulnerability*****:** Species where Concern Levels are expected to increase in the coming decades due to climate change.
    - ***Keystone Species*****:** Species that many other species rely on for their sustained presence.
    - ***Stronghold Species*****:** Species for which the Northeast supports the strongest populations and are imperiled outside of the region.
    - ***Genetic Distinctiveness*****:** Species or other taxonomic levels with unique genetics, such as isolated populations, DPS, subspecies, uncertain taxonomy, etc.
    - ***Cultural Values*****:** Species with historical significance or strong values to Indigenous peoples may be included as RSGCN in recognition of the importance of maintaining secure populations.
- Vertebrate and invertebrate taxa are screened with the same selection criteria.
- The Federal listing status criteria included Candidate species as well as Endangered, Threatened, or Proposed.
- The subnational rank or “S-Rank” filter is a regional average of all the states with an S-Rank for that species. A regional average S-Rank of less than S2 remains a filter. [used by Conservation Data Centres and NatureServe to rank the conservation status or rarity of species. Factors that are considered when ranking include the number of occurrences, population size, distribution, abundance trends, and threats. https://explorer.natureserve.org/AboutTheData/DataTypes/ConservationStatusCategories]
- A filter of State Protected Status (state regulatory listing) is now included for species prescreened as Maybe RSGCN.
- RSGCN Watchlist was added for species that are of concern to the taxa teams but for which:
    - The species are data deficient, have uncertain taxonomy, or are showing varying trends in different parts of the region, prioritizing them for additional survey or research efforts = **Watchlist [Assessment Priority]**
    - The species is interdependent with an RSGCN but does not qualify as RSGCN on its own = **Watchlist [Interdependent Species]**
    - The region has low regional responsibility but high concern = **Watchlist [Deferral to adjacent region]**
- Species that are not currently identified as SGCN by at least one state in the region may now be considered as **Proposed RSGCN** or **Proposed Watchlist** species.

Taxa teams remain the definitive authority on reviewing, confirming, or revising prescreened RSGCN recommendations, identifying Overriding Factor(s), determining RSGCN Concern Levels and Regional Responsibility, and recommending species for the Watchlist. Terwilliger Consulting Inc. coordinates their review and consensus process as part of the RCN Technical Services RCN project to the NEFWDTC. For more information on the methods and selection process see northeastwildlifediversity.org

### Appendix S3

#### Table of RSGCN Member Team (Current and Previous Members)

Table 1. Table of all contributions over time, anyone who has worked on the RSGCN lists formerly and the position they held within the NE while they contributed to the 2023 RSGCN list.

| Full Name | First Name | Last Name | Affiliation |
|---|---|---|---|
| Brad Allen | Brad | Allen | Maine Department of Inland Fisheries and Wildlife |
| Michael Amaral | Michael | Amaral | New Hampshire Fish and Game Department |
| Richard Bailey | Richard | Bailey | West Virginia Division of Natural Resources |
| Henrietta Bellman | Henrietta | Bellman | Delaware Department of Natural Resources and Environmental Control |
| Alyssa Bennett | Alyssa | Bennett | Vermont Fish and Wildlife Department |
| Jacqueline Benway | Jacqueline | Benway | Connecticut Department of Energy and Environmental Protection |
| Dee Blanton | Dee | Blanton | United States Fish and Wildlife Service |
| Brad Blodget | Brad | Blodget | Massachusetts Division of Fisheries and Wildlife |
| Ruth Boettcher | Ruth | Boettcher | Virginia Department of Wildlife Resources |
| Lisa Bonacci | Lisa | Bonacci | New York Department of Environmental Conservation |
| Robert Bourdon | Robert | Bourdon | National Oceanic and Atmospheric Administration |
| Daniel Bove | Daniel | Bove | Massachusetts Division of Fisheries and Wildlife |
| Jeanette Bowers | Jeanette | Bowers | New Jersey Department of Environmental Protection |

| | | | |
|---|---|---|---|
| Bethany Bradley | Bethany | Bradley | Northeast Climate Adaptation Science Center |
| Marc Branham | Marc | Branham | University of Florida at Gainesville |
| Dan Brauning | Dan | Brauning | Pennsylvania Game Commission |
| Al Breisch | Al | Breisch | New York Department of Environmental Conservation |
| Gwenda Brewer | Gwenda | Brewer | Maryland Department of Natural Resources |
| Charles Brown | Charles | Brown | Rhode Island Department of Environmental Management |
| Jeffrey Brust | Jeffrey | Brust | New Jersey Department of Environmental Protection |
| Scott Buchanan | Scott | Buchanan | Rhode Island Department of Environmental Management |
| David Bullis | David | Bullis | Syracuse University |
| Kevin Burgio | Kevin | Burgio | Northeast Climate Adaptation Science Center |
| Steve Burian | Steve | Burian | Southern Connecticut State University |
| Katharine Burns | Katharine | Burns | Rhode Island Department of Environmental Management |
| Scott Butterworth | Scott | Butterworth | West Virginia Division of Natural Resources |
| Jason Carmignani | Jason | Carmignani | Massachusetts Division of Fisheries and Wildlife |
| Matthew Carpenter | Matthew | Carpenter | New Hampshire Fish and Game Department |
| Casey Clark | Casey | Clark | Maine Department of Marine Resources |
| Kathy Clark | Kathy | Clark | New Jersey Department of Environmental Protection |
| Zack Couch | Zack | Couch | Office of Kentucky Nature Preserves |
| Elizabeth Crisfield | Elizabeth | Crisfield | Terwilliger Consulting Inc. |

| | | | |
|---|---|---|---|
| Shawn Crouse | Shawn | Crouse | New Jersey Department of Environmental Protection |
| Jason Davis | Jason | Davis | Delaware Department of Natural Resources and Environmental Control |
| Diana Day | Diana | Day | Pennsylvania Fish and Boat Commission |
| Phillip deMaynadier | Phillip | deMaynadier | Maine Department of Inland Fisheries and Wildlife |
| Paul Dennehy | Paul | Dennehy | Independent Lepidoptera expert |
| Randy Dettmers | Randy | Dettmers | United States Fish and Wildlife Service |
| Ed DeWalt | Ed | DeWalt | Illinois Natural History Survey |
| Jenny Dickson | Jenny | Dickson | Connecticut Department of Energy and Environmental Protection |
| Melissa Doperalski | Melissa | Doperalski | New Hampshire Fish and Game Department |
| Sam Droege | Sam | Droege | United States Geological Survey |
| Donovan Drummey | Donovan | Drummey | United States Fish and Wildlife Service |
| Kevin Eliason | Kevin | Eliason | West Virginia Division of Natural Resources |
| Julie Ellis | Julie | Ellis | University of Pennsylvania |
| Brian Eltz | Brian | Eltz | Connecticut Department of Energy and Environmental Protection |
| Claire Enterline | Claire | Enterline | Maine Department of Inland Fisheries and Wildlife |
| Candace Fallon | Candace | Fallon | Xerces Society |
| Lynn Faust | Lynn | Faust | Independent firefly expert |
| Frank Felbaum | Frank | Felbaum | Pennsylvania Department of Conservation and Natural Resources |
| Mark Ferguson | Mark | Ferguson | Vermont Fish and Wildlife Department |

| | | | |
|---|---|---|---|
| Doug Fischer | Doug | Fischer | Pennsylvania Fish and Boat Commission |
| Alexander Fish | Alexander | Fish | Maine Department of Inland Fisheries and Wildlife |
| Mack Frantz | Mack | Frantz | West Virginia Division of Natural Resources |
| Devaughn Fraser | Devaughn | Fraser | Connecticut Department of Energy and Environmental Protection |
| Amanda Freitas | Amanda | Freitas | Rhode Island Department of Environmental Management |
| Tom French | Tom | French | Massachusetts Division of Fisheries and Wildlife |
| Steve Fuller | Steve | Fuller | North Atlantic Landscape Conservation Cooperative |
| Merry Gallagher | Merry | Gallagher | Maine Department of Inland Fisheries and Wildlife |
| Catherine Gatenby | Catherine | Gatenby | United States Fish and Wildlife Service |
| Lisa Gelvin-Innvaer | Lisa | Gelvin-Innvaer | Connecticut Department of Energy and Environmental Protection |
| Kathy Gipe | Kathy | Gipe | Pennsylvania Fish and Boat Commission |
| Luke Groff | Luke | Groff | Vermont Fish and Wildlife Department |
| Becky Gwynn | Becky | Gwynn | Virginia Department of Wildlife Resources |
| Cathy Haffner | Cathy | Haffner | Pennsylvania Game Commission |
| Mackenzie Hall | Mackenzie | Hall | New Jersey Department of Environmental Protection |
| Sergio Harding | Sergio | Harding | Virginia Department of Wildlife Resources |
| Spencer Hardy | Spencer | Hardy | Vermont Center for Ecostudies |
| Jon Hare | Jon | Hare | National Oceanic and Atmospheric Administration |
| Alison Haskell | Alison | Haskell | United States Fish and Wildlife Service |

| | | | |
|---|---|---|---|
| Jerry Hassinger | Jerry | Hassinger | Pennsylvania Game Commission |
| Christopher Heckscher | Christopher | Heckscher | Delaware State University |
| John Heilferty | John | Heilferty | New Jersey Department of Environmental Protection |
| John Herbert | John | Herbert | Rhode Island Department of Environmental Management |
| Carl Herzog | Carl | Herzog | New York Department of Environmental Conservation |
| Brian Hess | Brian | Hess | Connecticut Department of Energy and Environmental Protection |
| Hanusia Higgins | Hanusia | Higgins | Northeast Climate Adaptation Science Center |
| Eric Hilton | Eric | Hilton | Virginia Institute of Marine Science |
| Heidi Holman | Heidi | Holman | New Hampshire Fish and Game Department |
| Lisa Holst | Lisa | Holst | New York Department of Environmental Conservation |
| Sandra Houghton | Sandra | Houghton | New Hampshire Fish and Game Department |
| Aaron Hunt | Aaron | Hunt | University of Delaware |
| Pam Hunt | Pam | Hunt | New Hampshire Audubon |
| Alan Hutchinson | Alan | Hutchinson | Maine Department of Inland Fisheries and Wildlife |
| Rob Jean | Rob | Jean | Environmental Solutions & Innovations Inc. |
| Scott Johnston | Scott | Johnston | United States Fish and Wildlife Service |
| Michael Jones | Michael | Jones | Massachusetts Division of Fisheries and Wildlife |
| Jennifer Kagel | Jennifer | Kagel | United States Fish and Wildlife Service |
| John Kanter | John | Kanter | New Hampshire Fish and Game Department |

| | | | |
|---|---|---|---|
| Jon Kart | Jon | Kart | Vermont Fish and Wildlife Department |
| Oliver Keller | Oliver | Keller | Florida Department of Agriculture |
| Greg Kenney | Greg | Kenney | New York Department of Environmental Conservation |
| Jay Kilian | Jay | Kilian | Maryland Department of Natural Resources |
| John Kleopfer | John | Kleopfer | Virginia Department of Wildlife Resources |
| Barry Knisley | Barry | Knisley | Randolph-Macon Colllege |
| Kathy Leo | Kathy | Leo | West Virginia Division of Natural Resources |
| Betsy Leppo | Betsy | Leppo | Western Pennsylvania Conservancy |
| Adrienne Leppold | Adrienne | Leppold | Maine Department of Inland Fisheries and Wildlife |
| Alan Libby | Alan | Libby | Rhode Island Department of Environmental Management |
| David Lieb | David | Lieb | Pennsylvania Fish and Boat Commission |
| Zachary Loughman | Zachary | Loughman | West Liberty University |
| Alice Lubeck | Alice | Lubeck | Northeast Climate Adaptation Science Center |
| Clare Maffei | Clare | Maffei | United States Fish and Wildlife Service |
| Amy Mahar | Amy | Mahar | New York Department of Environmental Conservation |
| John Maniscalco | John | Maniscalco | New York Department of Environmental Conservation |
| Michael Marchand | Michael | Marchand | New Hampshire Fish and Game Department |
| Amy Martin | Amy | Martin | Virginia Department of Wildlife Resources |
| Larry Master | Larry | Master | The Nature Conservancy |

| | | | |
|---|---|---|---|
| Jonathan Mawdsley | Jonathan | Mawdsley | United States Geological Survey |
| James McCann | James | McCann | Maryland Department of Natural Resources |
| Max McCarthy | Max | McCarthy | Rutgers University |
| Mark McCullough | Mark | McCullough | Maine Department of Inland Fisheries and Wildlife |
| Chris McDowell | Chris | McDowell | Connecticut Department of Energy and Environmental Protection |
| Mathew McKinney | Mathew | McKinney | West Virginia Division of Natural Resources |
| Jonathan McKnight | Jonathan | McKnight | Maryland Department of Natural Resources |
| Conor McManus | Conor | McManus | Rhode Island Department of Environmental Management |
| Dawn McReynolds | Dawn | McReynolds | New York Department of Environmental Conservation |
| Mark Mello | Mark | Mello | Lloyd Center for the Environment |
| Scott Melvin | Scott | Melvin | Massachusetts Division of Fisheries and Wildlife |
| Deb Mignogna | Deb | Mignogna | United States Fish and Wildlife Service |
| Joan Milam | Joan | Milam | University of Massachusetts Amherst |
| Robert Miller | Robert | Miller | New York Department of Environmental Conservation |
| Jorge Montero | Jorge | Montero | Anacostia Watershed Society |
| Johnny Moore | Johnny | Moore | Delaware Department of Natural Resources and Environmental Control |
| Clinton Morgeson | Clinton | Morgeson | Virginia Department of Wildlife Resources |
| Doug Morin | Doug | Morin | Vermont Fish and Wildlife Department |
| Margaret Murphy | Margaret | Murphy | Vermont Fish and Wildlife Department |

| | | | |
|---|---|---|---|
| Sean Murphy | Sean | Murphy | Pennsylvania Game Commission |
| Luke Myers | Luke | Myers | State University of New York Plattsburgh |
| Nathan Nazdrowicz | Nathan | Nazdrowicz | Delaware Department of Natural Resources and Environmental Control |
| Alicia Nelson | Alicia | Nelson | Virginia Marine Resources Commission |
| Michael Nelson | Michael | Nelson | Massachusetts Division of Fisheries and Wildlife |
| Holly Niederriter | Holly | Niederriter | Delaware Department of Natural Resources and Environmental Control |
| Larry Niles | Larry | Niles | New Jersey Department of Environmental Protection |
| Peter Nye | Peter | Nye | New York Department of Environmental Conservation |
| Kathleen O'Brien | Kathleen | O'Brien | New York Department of Environmental Conservation |
| Susan Olcott | Susan | Olcott | West Virginia Division of Natural Resources |
| Scott Olszewski | Scott | Olszewski | Rhode Island Department of Environmental Management |
| Kieran O'Malley | Kieran | O'Malley | West Virginia Division of Natural Resources |
| Richard Orr | Richard | Orr | Mid-Atlantic Invertebrate Field Studies |
| Damien Ossi | Damien | Ossi | District of Columbia Department of Energy and Environment |
| Nathaniel Owens | Nathaniel | Owens | West Virginia Division of Natural Resources |
| Kevin Oxenreider | Kevin | Oxenreider | West Virginia Division of Natural Resources |
| Matthew Palumbo | Matthew | Palumbo | New York Department of Environmental Conservation |
| Steve Parren | Steve | Parren | Vermont Fish and Wildlife Department |
| Timothy Pearce | Timothy | Pearce | Carnegie Museum of Natural History |

| | | | |
|---|---|---|---|
| Corey Pelletier | Corey | Pelletier | Rhode Island Department of Environmental Management |
| Diane Pence | Diane | Pence | United States Fish and Wildlife Service |
| Mike Pinder | Mike | Pinder | Virginia Department of Wildlife Resources |
| Joe Racette | Joe | Racette | New York Department of Environmental Conservation |
| Chris Raithel | Chris | Raithel | Rhode Island Department of Environmental Management |
| Valinn Ranelli | Valinn | Ranelli | Independent Odonata expert |
| Andrew Rasmussen | Andrew | Rasmussen | Florida A & M University |
| Daniel Rauch | Daniel | Rauch | District of Columbia Department of Energy and Environment |
| Michael Ravesi | Michael | Ravesi | Connecticut Department of Energy and Environmental Protection |
| Rosalind Renfrew | Rosalind | Renfrew | Vermont Fish and Wildlife Department |
| Rick Reynolds | Rick | Reynolds | Virginia Department of Wildlife Resources |
| Tracy Monegan Rice | Tracy Monegan | Rice | Terwilliger Consulting Inc. |
| Leif Richardson | Leif | Richardson | Xerces Society |
| Calvin Ritter | Calvin | Ritter | United States Fish and Wildlife Service |
| Samantha Robinson | Samantha | Robinson | Delaware Department of Natural Resources and Environmental Control |
| Katherine Rodrigue | Katherine | Rodrigue | Rhode Island Department of Environmental Management |
| Teresa Rodriguez | Teresa | Rodriguez | District of Columbia Department of Energy and Environment |
| Joseph Rogerson | Joseph | Rogerson | Delaware Department of Natural Resources and Environmental Control |
| Lindsey Rohrbaugh | Lindsey | Rohrbaugh | District of Columbia Department of Energy and Environment |

| | | | |
|---|---|---|---|
| Dan Rosenblatt | Dan | Rosenblatt | New York Department of Environmental Conservation |
| Daniel Ryan | Daniel | Ryan | District of Columbia Department of Energy and Environment |
| Natalie Sacco | Natalie | Sacco | New York Department of Environmental Conservation |
| Laura Saucier | Laura | Saucier | Connecticut Department of Energy and Environmental Protection |
| Kris Schantz | Kris | Schantz | New Jersey Department of Environmental Protection |
| Eve Schluter | Eve | Schluter | Massachusetts Division of Fisheries and Wildlife |
| Dale Schweitzer | Dale | Schweitzer | Connecticut Department of Energy and Environmental Protection |
| Jennifer Selfridge | Jennifer | Selfridge | Maryland Department of Natural Resources |
| Virginia Shepherd | Virginia | Shepherd | Terwilliger Consulting Inc. |
| Andrew Shiels | Andrew | Shiels | Pennsylvania Fish and Boat Commission |
| Julie Slacum | Julie | Slacum | United States Fish and Wildlife Service |
| Scott Smith | Scott | Smith | Maryland Department of Natural Resources |
| Robert Somes | Robert | Somes | New Jersey Department of Environmental Protection |
| Caleb Spiegel | Caleb | Spiegel | United States Fish and Wildlife Service |
| Michael Stangl | Michael | Stangl | Delaware Department of Natural Resources and Environmental Control |
| Melissa Starking | Melissa | Starking | Terwilliger Consulting Inc. |
| Michelle Staudinger | Michelle | Staudinger | Northeast Climate Adaptation Science Center |
| Cory Stearns | Cory | Stearns | Maine Department of Inland Fisheries and Wildlife |
| Chuck Stence | Chuck | Stence | Maryland Department of Natural Resources |

| | | | |
|---|---|---|---|
| Edna Stetzar | Edna | Stetzar | Delaware Department of Natural Resources and Environmental Control |
| Craig Stihler | Craig | Stihler | West Virginia Division of Natural Resources |
| David Stormer | David | Stormer | Delaware Department of Natural Resources and Environmental Control |
| Beth Swartz | Beth | Swartz | Maine Department of Inland Fisheries and Wildlife |
| Jordan Terrell | Jordan | Terrell | Delaware Department of Natural Resources and Environmental Control |
| Karen Terwilliger | Karen | Terwilliger | Terwilliger Consulting Inc. |
| Glen Therres | Glen | Therres | Maryland Department of Natural Resources |
| Roger Thoma | Roger | Thoma | The Ohio State University |
| Jeff Trollinger | Jeff | Trollinger | Virginia Department of Wildlife Resources |
| Anthony Tur | Anthony | Tur | United States Fish and Wildlife Service |
| Maria Tur | Maria | Tur | United States Fish and Wildlife Service |
| Greg Turner | Greg | Turner | Pennsylvania Game Commission |
| Chris Urban | Chris | Urban | Pennsylvania Fish and Boat Commission |
| Rick Van de Poll | Rick | Van de Poll | Independent Lepidoptera expert |
| Julie Victoria | Julie | Victoria | Connecticut Department of Energy and Environmental Protection |
| Andrew Vitz | Andrew | Vitz | Massachusetts Division of Fisheries and Wildlife |
| Susie von Oettingen | Susie | von Oettingen | United States Fish and Wildlife Service |
| David Wagner | David | Wagner | University of Connecticut |
| Jim Wagner | Jim | Wagner | Western Pennsylvania Conservancy |

| | | | |
|---|---|---|---|
| Steve Walker | Steve | Walker | Maine Department of Inland Fisheries and Wildlife |
| Mary Walsh | Mary | Walsh | Western Pennsylvania Conservancy |
| Wendy Walsh | Wendy | Walsh | United States Fish and Wildlife Service |
| Brian Watson | Brian | Watson | Virginia Department of Wildlife Resources |
| Nevin Welte | Nevin | Welte | Pennsylvania Fish and Boat Commission |
| Angel Willey | Angel | Willey | Maryland Department of Natural Resources |
| Lisa Williams | Lisa | Williams | Pennsylvania Game Commission |
| Alissa Wilson | Alissa | Wilson | New Jersey Department of Environmental Protection |
| Rachael Winfree | Rachael | Winfree | Rutgers University |
| Pete Woods | Pete | Woods | Western Pennsylvania Conservancy |
| Derek Yorks | Derek | Yorks | Maine Department of Inland Fisheries and Wildlife |
| Brian Zarate | Brian | Zarate | New Jersey Department of Environmental Protection |
| Tracy Zarrillo | Tracy | Zarrillo | Connecticut Department of Energy and Environmental Protection |

### Appendix S4

#### Tables of Data Sources (including all taxonomic authorities)

Table 5-1 of online sources for Pre-screening data.

| Name | Type of Information | URL |
|---|---|---|
| Integrated Taxonomic Information System (ITIS) | taxonomy | https://www.itis.gov/ |
| IUCN Redlist of Threatened Species | distribution, ecology, threats, conservation ranks | https://www.iucnredlist.org/ |
| NatureServe | distribution, ecology, threats, conservation ranks | https://explorer.natureserve.org/ |
| Botanical Information and Ecology Network (BIEN) | distribution, ecology | https://bien.nceas.ucsb.edu/bien/ |
| USFWS National Listing Workplan | conservation ranks | https://www.fws.gov/project/national-listing-workplan |
| Environmental Conservation Online System (ECOS) | conservation ranks | https://ecos.fws.gov/ecp/ |
| Ocean Biodiversity Information System (OBIS) | distribution | https://obis.org/ |

Table 5-2 of Taxonomic Authorities

| Name | Taxonomic Group | Type | URL |
|---|---|---|---|
| Integrated Taxonomic Information System (ITIS) | all | online | https://www.itis.gov/ |
| American Ornithological Union/Society (AOU/S) checklist of north and middle American birds | birds | online | https://checklist.americanornithology.org/ |
| American Birding Association (ABA) checklist | birds | online | https://www.aba.org/listing-taxonomy/ |

| | | | |
|---|---|---|---|
| An updated classification of the freshwater crayfishes of the world, with a complete species list | crayfish | journal article | https://academic.oup.com/jcb/article/37/5/615/4060680 |
| Ephemeroptera of the world | ephemeroptera | online | https://insecta.bio.spbu.ru/z/Eph-spp/ |
| Firefly species checklist of the USA and Canada | fireflies | online | https://www.fireflyatlas.org/firefly-species/firefly-species-checklist |
| A naturalist's long walk among the shadows: of North American *Photuris* - patterns, outlines, silhouettes … echoes | fireflies | book | https://bugguide.net/node/view/1533445 |
| Merican Fisheries Society (AFS) | fishes | onlne | https://fisheries.org/ |
| Eschmeyer's catalog of fishes online database | fishes | online | https://www.calacademy.org/scientists/projects/eschmeyers-catalog-of-fishes |
| Fishes of the world, 5th edition | fishes | book | https://www.wiley.com/en-us/Fishes+of+the+World%2C+5th+Edition-p-9781119220824 *Free version of the 4th edition*: http://www.sisal.unam.mx/labeco/LAB_ECOLOGIA/Ecologia_de_peces_files/Nelson%202006.pdf |
| Common and scientific names of fishes from the United States, Canada, and Mexico, 8th edition | fishes | book | https://fisheries.org/bookstore/all-titles/special-publications/namesoffishes8/ *Free version of the 7th edition*: https://downloads.regulations.gov/FWS-R1-ES-2017-0035-0004/attachment_23.pdf |
| A revised list of the freshwater mussels (Mollusca: Bivalvia: Unionida) of the United States and Canada | freshwater mussels | journal article | https://doi-org.fwslibrary.idm.oclc.org/10.31931/fmbc.v20i2.2017.33-58 |
| Scientific and standard English names of amphibians and reptiles of North America north of Mexico, 8th edition | herptiles | book | https://ssarherps.org/publications/north-american-checklist/ |
| Discover Life bee species guide and world checklist | hymenoptera | online | https://www.discoverlife.org/mp/20q?act=x_checklist&guide=Apoidea_species |
| A catalogue of the butterflies of the United States and Canada | lepidoptera | online | https://butterfliesofamerica.com/US-Can-Cat.htm |
| Illustrated lists of American butterflies (North and South America) | lepidoptera | online | https://www.butterfliesofamerica.com/L/Neotropical.htm |
| Mammal diversity database | mammals | online | https://www.mammaldiversity.org/ |

| | | | |
|---|---|---|---|
| World Register of Marine Species (WoRMS) | marine species | online | https://www.marinespecies.org/index.php |
| Molluscabase | molluscs | online | https://www.molluscabase.org/about.php |
| World odonata list: OdonataCentral | odonata | online | https://www.odonatacentral.org/app/#/wol/ |
| Plecoptera species file | plecoptera | online | http://plecoptera.archive.speciesfile.org/HomePage/Plecoptera/HomePage.aspx |
| Trichoptera world checklist database | trichoptera | online | https://trichopt.app.clemson.edu/welcome.php |
| Trichoptera Nearctica | trichoptera | online | https://trichoptera.org/ |

Table 5-3 State Data Sources for pre-screening.

| State | Type of Information | Wildlife (Mammals, Birds, Herptiles) | Fish | Aquatic Inverts (mussels, crayfish, snails, sponges, etc.) | Terrestrial Inverts (millepedes, insects, arachnids, snails, etc.) | Plants | Marine Mammals/ Sea Turtles | Marine Fish | Marine Inverts | URL |
|---|---|---|---|---|---|---|---|---|---|---|
| CT | SGCN status | Yes | Yes | Yes | Yes | Yes | Yes | Yes | Yes | https://portal.ct.gov/deep/wildlife/ct-wildlife-action-plan/ct-2015-wildlife-action-plan |
| DC | SGCN status | Yes | Yes | Yes | Yes | | | | | https://doee.dc.gov/service/wildlifeactionplan |
| DE | SGCN status | Yes | Yes | Yes | Yes | | Yes | Yes | Yes | https://dnrec.delaware.gov/fish-wildlife/conservation/wildlife-action-plan/ |
| MA | SGCN status | Yes | Yes | Yes | Yes | Yes | Yes | | | https://www.mass.gov/info-details/state-wildlife-action-plan-swap |
| MD | SGCN status | Yes | Yes | Yes | Yes | Yes | Yes | Yes | Yes | https://dnr.maryland.gov/wildlife/Pages/plants_wildlife/SWAP_Submission.aspx |
| ME | SGCN status | Yes | Yes | Yes | Yes | | Yes | Yes | Yes | https://www.maine.gov/ifw/fish-wildlife/wildlife/wildlife-action-plan/index.html |

| | | | | | | | | | | |
|---|---|---|---|---|---|---|---|---|---|---|
| NH | SGCN status | Yes | Yes | Yes | Yes | | Yes | | Yes | https://www.wildlife.nh.gov/wildlife-and-habitat/nh-wildlife-action-plan/swap-2015 |
| NJ | SGCN status | Yes | Yes | Yes | Yes | | Yes | Yes | Yes | https://dep.nj.gov/njfw/wildlife/new-jerseys-state-wildlife-action-plan/ |
| NY | SGCN status | Yes | Yes | Yes | Yes | | Yes | Yes | Yes | https://dec.ny.gov/nature/animals-fish-plants/biodiversity-species-conservation/state-wildlife-action-plan |
| PA | SGCN status | Yes | Yes | Yes | Yes | | | | | https://www.pgc.pa.gov/Wildlife/WildlifeActionPlan/Pages/default.aspx |
| RI | SGCN status | Yes | Yes | Yes | Yes | Yes | Yes | Yes | Yes | https://dem.ri.gov/natural-resources-bureau/fish-wildlife/wildlife-hunting/ri-state-wildlife-action-plan |
| VA | SGCN status | Yes | Yes | Yes | Yes | | Yes | | | https://dwr.virginia.gov/wildlife/wildlife-action-plan/wildlife-action-plan-2015/ |
| VT | SGCN status | Yes | Yes | Yes | Yes | Yes | n/a | n/a | n/a | https://vtfishandwildlife.com/about-us/budget-and-planning/wildlife-action-plan |
| WV | SGCN status | Yes | Yes | Yes | Yes | Yes | n/a | n/a | n/a | https://wvdnr.gov/state-wildlife-action-plan/ |
| CT | conservation status | Yes | Yes | Yes | Yes | Yes | Yes | Yes | Yes | https://portal.ct.gov/deep/endangered-species/connecticuts-endangered-threatened-and-special-concern-species |
| DC | conservation status | | | | | | | | | *see below |
| DE | conservation status | Yes | Yes | Yes | Yes | | | | | https://dnrec.delaware.gov/fish-wildlife/conservation/endangered-species/ |
| MA | conservation status | Yes | Yes | Yes | Yes | Yes | Yes | | | https://www.mass.gov/info-details/list-of-endangered-threatened-and-special-concern-species |
| MD | conservation status | Yes | Yes | Yes | Yes | Yes | Yes | Yes | Yes | https://dnr.maryland.gov/wildlife/Pages/plants_wildlife/rte/rteanimals.aspx<br>https://dnr.maryland.gov/wildlife/pages/plants_wildlife/rte/rteplants.aspx<br>https://dnr.maryland.gov/fisheries/Pages/endangered.aspx |
| ME | conservation status | Yes | Yes | Yes | Yes | Yes | Yes | Yes | | https://www.maine.gov/ifw/fish-wildlife/wildlife/endangered-threatened-species/listed-species.html<br>https://www.maine.gov/ifw/fish-wildlife/wildlife/endangered-threatened-species/special-concern.html<br>https://www.mainelegislature.org/legis/statutes/12/title12sec697 |

| | | | | | | | | | | |
|---|---|---|---|---|---|---|---|---|---|---|
| | | | | | | | | | | 5.html https://www.maine.gov/dacf/mnap/features/rare_plants/index.htm |
| NH | conservation status | Yes | Yes | Yes | Yes | Yes | | | | https://www.wildlife.nh.gov/wildlife-and-habitat/nongame-and-endangered-species/endangered-and-threatened-wildlife-nh https://www.nhdfl.dncr.nh.gov/natural-heritage/rare-native-plants |
| NJ | conservation status | Yes | Yes | Yes | Yes | Yes | Yes | | | https://dep.nj.gov/njfw/wildlife/endangered-threatened-and-special-concern-species/ https://nj.gov/dep/parksandforests/natural/docs/njplantlist.pdf |
| NY | conservation status | Yes | Yes | Yes | Yes | Yes | Yes | | | https://dec.ny.gov/nature/animals-fish-plants/biodiversity-species-conservation/endangered-species/list https://dec.ny.gov/nature/animals-fish-plants/plants/state-protected-plants |
| PA | conservation status | Yes | Yes | Yes | | Yes | | | | https://www.pgc.pa.gov/Wildlife/EndangeredandThreatened/Pages/default.aspx https://www.fishandboat.com/Conservation/Threatened-and-Endangered-Species/Pages/default.aspx https://www.dcnr.pa.gov/Conservation/WildPlants/RareThreatenedAndEndangeredPlants/Pages/default.aspx |
| RI | conservation status | Yes | Yes | Yes | Yes | Yes | Yes | | | https://rinhs.org/species/rare-species/ |
| VA | conservation status | Yes | Yes | Yes | Yes | Yes | Yes | | | https://dwr.virginia.gov/wp-content/uploads/media/virginia-threatened-endangered-species.pdf https://www.vdacs.virginia.gov/plant-industry-services-endangered-species.shtml |
| VT | conservation status | Yes | Yes | Yes | Yes | Yes | n/a | n/a | n/a | https://vtfishandwildlife.com/conserve/endangered-and-threatened-species |
| WV | conservation status | | | | | | n/a | n/a | n/a | *natural heritage programs - email Damien and Sophie for DC, K for WV |

### Appendix S5

#### Taxonomic Team Sheet Example

Shared as an excel workbook
